# Tumor elimination by clustered microRNAs miR-306 and miR-79 via non-canonical activation of JNK signaling

**DOI:** 10.1101/2022.02.11.480121

**Authors:** Zhaowei Wang, Xiaoling Xia, Tatsushi Igaki

**Affiliations:** State Key Laboratory of Biocontrol, School of Ecology, Sun Yat-sen University, Shenzhen, Guangdong, 518107, China; Laboratory of Genetics, Graduate School of Biostudies, Kyoto University, Yoshida-Konoe-cho, Sakyoku, Kyoto, 607-8501, Japan; Guangzhou Key Laboratory of Insect Development Regulation and Application Research, Institute of Insect Science and Technology & School of Life Sciences, South China Normal University, Guangzhou, Guangdong, 510631, China

## Abstract

JNK signaling plays a critical role in both tumor promotion and tumor suppression. Here, we identified clustered microRNAs (miRNAs) miR-306 and miR-79 as novel tumor-suppressor miRNAs that specifically eliminate JNK-activated tumors in *Drosophila*. While showing no significant effect on normal tissue growth, miR-306 and miR-79 strongly suppressed growth of multiple tumor models including malignant tumors caused by Ras activation and cell polarity defects. Mechanistically, these miRNAs commonly target the mRNA of an E3 ubiquitin ligase *Drosophila* ring finger protein 146 (dRNF146). We found that DRNF146 promotes degradation of tankyrase (Tnks), an ADP-ribose polymerase that promotes JNK activation in a non-canonical manner. Thus, downregulation of dRNF146 by miR-306 and miR-79 leads to hyper-enhancement of JNK activation. Our data show that, while JNK activity is essential for tumor growth, elevation of miR-306 or miR-79 overactivate JNK signaling to the lethal level via non-canonical JNK pathway and thus eliminate tumors, providing a new miRNA-based strategy against cancer.

## Introduction

Cancer progression is driven by oncogenic alterations of intracellular signaling that lead to promotion of cell proliferation and suppression of cell death (Croce, 2008). The c-Jun N-terminal kinase (JNK) pathway is an evolutionarily conserved mitogen-activated protein (MAP) kinase cascade that regulates both cell proliferation and cell death in normal development and cancer (Bode & Dong, 2007; Eferl & Wagner, 2003). Indeed, JNK signaling can act as both tumor promoter and tumor suppressor depending on the cellular contexts (Bode & Dong, 2007; Bubici & Papa, 2014; Karin & Gallagher, 2005). Crucially, JNK signaling is often activated in various types of cancers (Bubici & Papa, 2014; Wu et al., 2019). Thus, accumulating evidence suggests that JNK signaling can be a critical therapeutic target for cancer. For instance, converting JNK’s role from pro-tumor to anti-tumor within tumor tissue could be an ideal anti-cancer strategy.

*Drosophila* provides a superb model for studying the genetic pathway of cellular signaling and has made great contributions to understand the basic principle of tumor growth and progression (Enomoto, Siow, & Igaki, 2018; Tipping & Perrimon, 2014). The best-studied model of *Drosophila* malignant tumor is generated by clones of cells overexpressing oncogenic Ras (Ras^V12^) with simultaneous mutations in apicobasal polarity genes such as *lethal giant larvae* (*lgl*), *scribble* (*scrib*) or *discs large* (*dlg*) in the imaginal epithelium (Brumby & Richardson, 2003; Pagliarini & Xu, 2003). These tumors activate JNK signaling and blocking JNK within the clones strongly suppresses their tumor growth (Igaki, Pagliarini, & Xu, 2006; Uhlirova & Bohmann, 2006), indicating that JNK acts as a pro-tumor signaling in these malignant tumors. Conversely, clones of cells overexpressing the oncogene Src in the imaginal discs activate JNK signaling and blocking JNK in these clones results in an enhanced overgrowth (Enomoto & Igaki, 2013), indicating that JNK negatively regulates Src-induced tumor growth. Similarly, although clones of cells mutant for *scrib* or *dlg* in the imaginal discs are eliminated by apoptosis when surrounded by wild-type cells, blocking JNK in these clones suppress elimination and causes tumorous overgrowth (Brumby & Richardson, 2003; Igaki, Pastor-Pareja, Aonuma, Miura, & Xu, 2009), indicating that JNK acts as anti-tumor signaling in these mutant clones. Thus, JNK also acts as both pro- and anti-tumor signaling depending on the cellular contexts in *Drosophila* imaginal epithelium.

miRNAs are a group of small non-coding RNAs that suppress target gene expression by mRNA degradation or translational repression and have been proposed to be potent targets for cancer therapy. Indeed, several cancer-targeted miRNA drugs have entered clinical trials in recent years. For instance, MRX34, a miRNA mimic drug developed from the tumor suppressor miR-34a, is the first miRNA-based anti-cancer drug that have entered Phase I clinical trials for patients with advanced solid tumors (Beg et al., 2017; Hong et al., 2020). In addition, MesomiR-1, a miR-16 mimic miRNA that targets EGFR, has entered Phase I trial for the treatment of thoracic cancers (Reid et al., 2013; van Zandwijk et al., 2017). Such miRNA-mediated anti-cancer strategy can be studied using the *Drosophila* tumor models. Indeed, in *Drosophila*, the conserved miRNA let-7 targets a transcription factor *chinmo* and thus suppresses tumor growth caused by *polyhomeotic* mutations (Jiang, Seimiya, Schlumpf, & Paro, 2018). In addition, miR-8 acts as a tumor suppressor against Notch-induced *Drosophila* tumors by directly inhibiting the Notch ligand Serrate (Vallejo, Caparros, & Dominguez, 2011). However, apart from these miRNAs that suppress growth of specific types of tumors, it is unclear whether there exist miRNAs that generally suppress tumor growth caused by different genetic alterations.

Here, using *Drosophila* tumor models and subsequent genetic analyses, we identified several tumor-suppressor miRNAs. Among these, miR-306 and miR-79, two clustered miRNAs located on the miR-9c/306/79/9b cluster, significantly suppressed growth of multiple types of JNK-activated tumors while showing only a slight effect on normal tissue growth. Mechanistically, miR-306 and miR-79 directly target *dRNF146*, an E3 ubiquitin ligase that causes degradation of a JNK-promoting ADP-ribose polymerase Tnks, thereby over-amplifying JNK signaling in tumors to the lethal levels via non-canonical JNK activation. Our findings provide a novel miRNA-based strategy that generally suppress growth of JNK-activating tumors.

## Results

### Identification of miR-306 and miR-79 as novel tumor-suppressor miRNAs

To identify novel anti-tumor miRNAs in *Drosophila*, we focused on 37 miRNA clusters or miRNAs that are highly expressed in *Drosophila* eye-antennal discs (Chung, Okamura, Martin, & Lai, 2008). Using the Flippase (FLP)-Flp recognition target (FRT)-mediated genetic mosaic technique, each miRNA was overexpressed in clones of cells expressing Ras^V12^ with simultaneous mutations in the apicobasal polarity gene *dlg* (Ras^V12^/*dlg^−/−^*) in the eye-antennal discs, the best-studied malignant tumor model in *Drosophila* (Pagliarini & Xu, 2003) (Figure 1A, compare to Figure 1K). We found that overexpression of miR-7, miR-79, miR-252, miR-276a, miR-276b, miR-282, miR-306, miR-310, miR-317, miR-981, miR-988, or the miR-9c/306/79/9b cluster in Ras^V12^/*dlg^−/−^* clones dramatically suppressed tumor growth (Figure 1A-E, Figure 1—figure supplement 1D, R, S, T, W, Y, Z, AC, and AE, quantified in Figure 1F and Figure 1—figure supplement 1AI). In addition, overexpression of miR-305, miR-995, or the miR-13a/13b-1/2c cluster mildly suppressed Ras^V12^/*dlg^−/−^* tumor growth (Figure 1—figure supplement 1K, X and AF, quantified in Figure 1—figure supplement 1AI). Clustered miRNAs are localized close to each other in the genome and are thus normally transcribed together, ensuring the transcription efficiency of miRNA genes (Kabekkodu et al., 2018; Ryazansky, Gvozdev, & Berezikov, 2011). Notably, overexpression of the miR-9c/306/79/9b cluster, miR-306, or miR-79 dramatically inhibited Ras^V12^/*dlg^−/−^* tumor growth (Figure 1C-E, compare to Figure 1B, quantified in Figure 1F). In addition, overexpression of miR-306 or miR-79 was sufficient to rescue the reduced pupation rate and animal lethality caused by Ras^V12^/*dlg^−/−^* tumors in the eye-antennal discs (Figure 1G-J). A pervious study in *Drosophila* wing discs showed that overexpression of miR-79 suppressed tumor growth caused by coexpression of Ras^V12^ and *lgl*-RNAi via unknown mechanisms (Shu et al., 2017). Similarly, we found that overexpression of the miR-9c/306/79/9b cluster, miR-306, or miR-79 strongly suppressed growth of Ras^V12^/*lgl^−/−^* tumors (Figure 1—figure supplement 2A-D, quantified in Figure 1—figure supplement 2E). Importantly, overexpression of the miR-9c/306/79/9b cluster, miR-306, or miR-79 alone only slightly reduced clone size compared to wild-type (Figure 1K-N, quantified in Figure 1O). These data indicate that miR-306 and miR-79 are tumor-suppressor miRNAs that only mildly suppress normal tissue growth but specifically blocks tumor growth in *Drosophila* imaginal epithelium.

**Figure 1.**
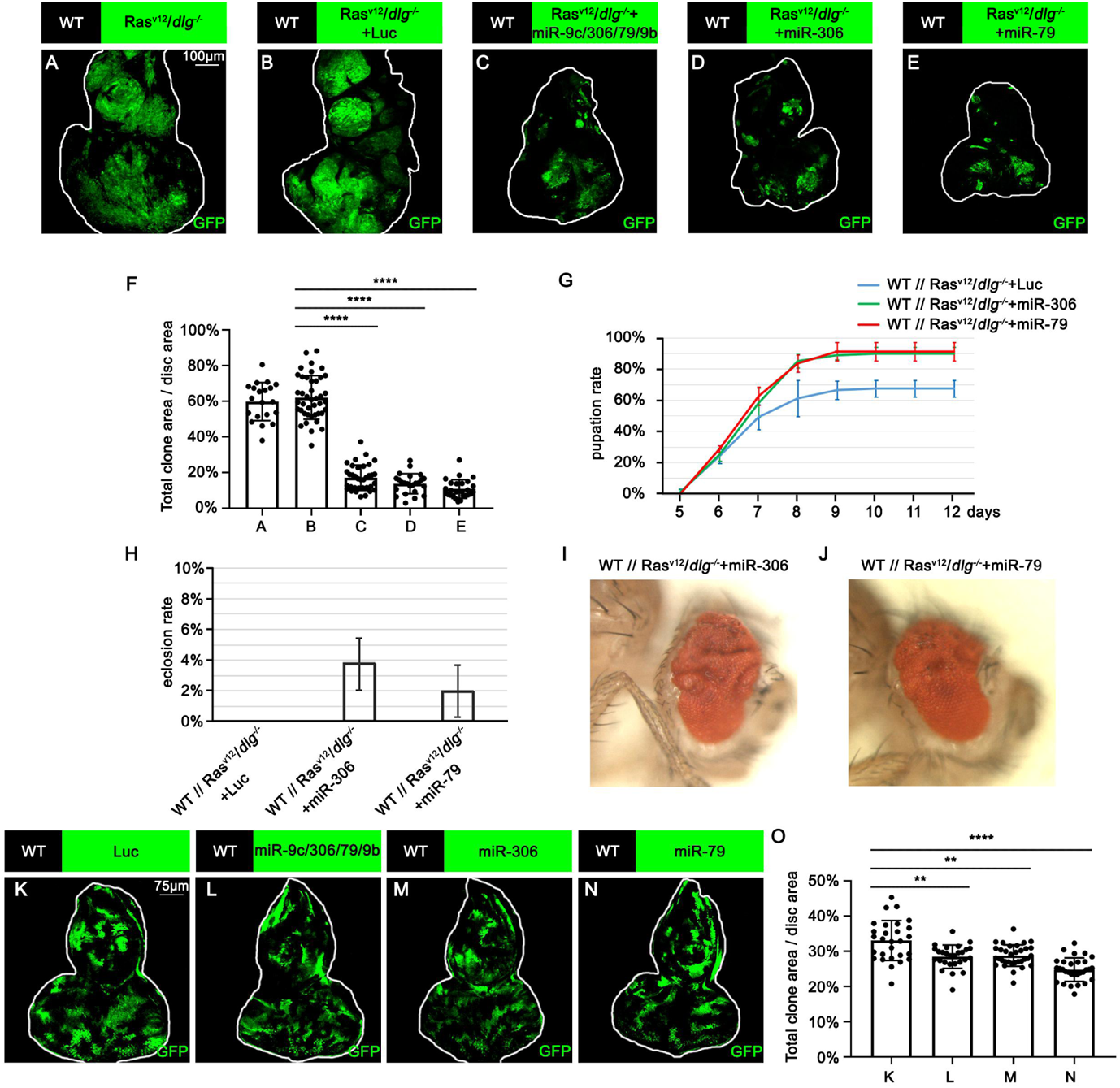
miR-306 and miR-79 suppress Ras^V12^/*dlg^−/−^* tumor growth. (A-E) Eye-antennal disc bearing GFP-labeled clones of indicated genotypes (7 days after egg laying). (F) Quantification of clone size (% of total clone area per disc area in eye-antennal disc) in (A-E). Error bars, SD; ****, p<0.0001 by one-way ANOVA multiple comparison test. (G) Pupation rate of flies with indicated genotypes. Data from three independent experiment, n>30 for each group in one experiment; error bars, SD. Eclosion rate of flies with indicated genotypes. Data from three independent experiment, n>30 for each group in one experiment; error bars, SD. (I-J) Adult eye phenotype of flies with indicated genotypes. (K-N) Eye-antennal disc bearing GFP-labeled clones of indicated genotypes (5 days after egg laying). (O) Quantification of clone size (% of total clone area per disc area in eye-antennal disc) of (K-N). Error bars, SD;**, p<0.01, ****, p<0.0001 by one-way ANOVA multiple comparison test.

### miR-306 and miR-79 suppress tumor growth by promoting cell death

We next investigated the mechanism by which miR-306 and miR-79 suppress tumor growth. Immunostaining of Ras^V12^/*dlg^−/−^* or Ras^V12^/*lgl^−/−^* tumors with anti-cleaved DCP-1 antibody revealed that expression of the miR-9c/306/79/9b cluster, miR-306, or miR-79 in tumor clones significantly increased the number of dying cells (Figure 2A-D, Figure 2—figure supplement 1A-D, quantified in Figure 2E and Figure 2—figure supplement 1E). In addition, blocking cell death in tumor clones by overexpressing the caspase inhibitor baculovirus p35 cancelled the tumor-suppressive activity of miR-306 or miR-79, while p35 overexpression alone did not affect growth of Ras^V12^/*dlg^−/−^* tumors (Figure 2F-H, quantified in Figure 2I). These data indicate that the miR-9c/306/79/9b cluster, miR-306, or miR-79 suppresses tumor growth by inducing cell death. Importantly, overexpression of these miRNAs alone did not cause cell death in normal tissue (Figure 2J-M, quantified in Figure 2N), suggesting that miR-306 or miR-79 cooperates with a putative tumor-specific signaling activated in Ras^V12^/*dlg^−/−^* or Ras^V12^/*lgl^−/−^* tumors to induce synthetic lethality.

**Figure 2.**
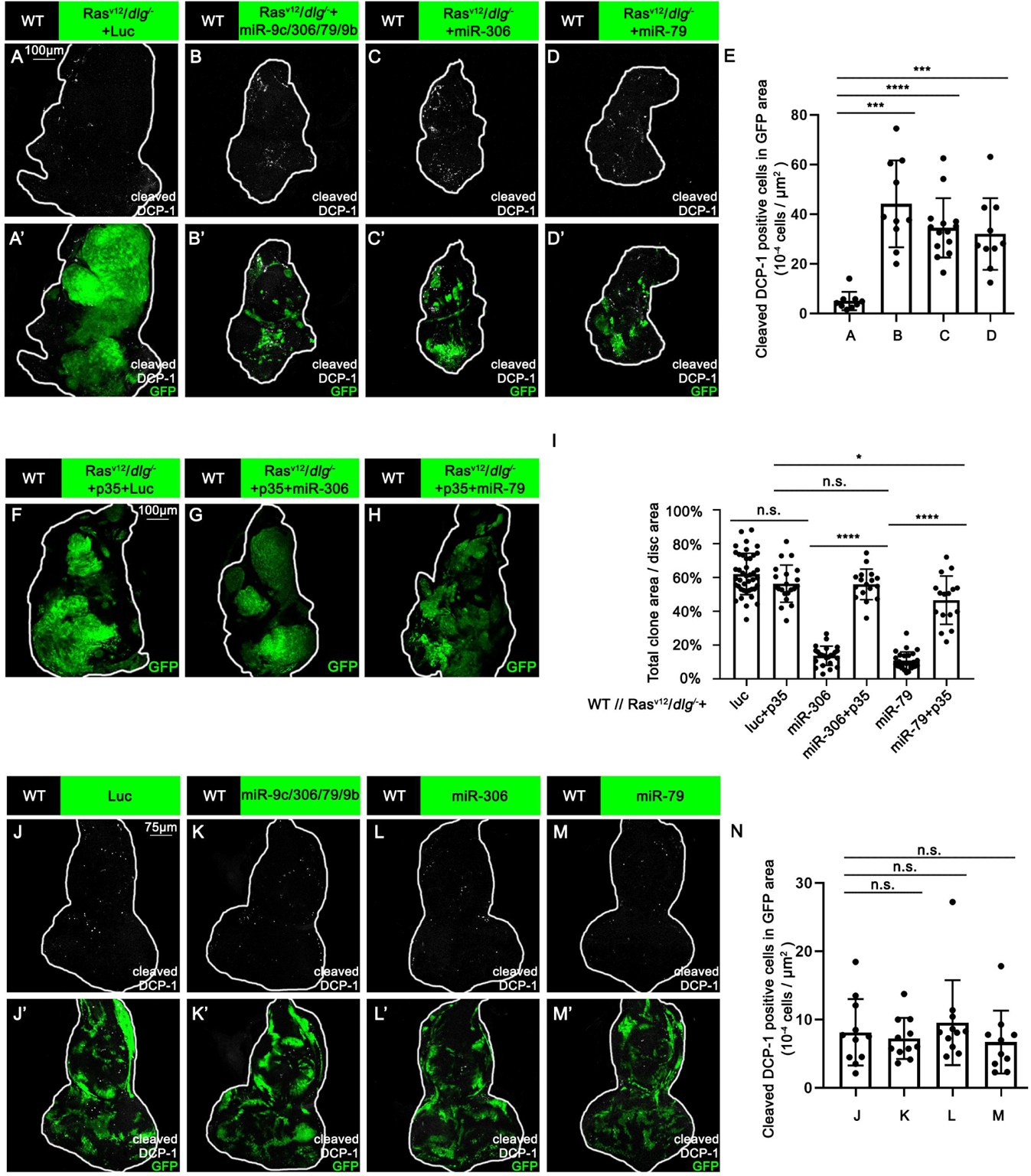
miR-306 and mir-79 suppress Ras^V12^*/dlg^−/−^* tumor growth by inducing apoptosis. (A-D) Eye-antennal disc bearing GFP-labeled clones of indicated genotypes stained with anti-cleaved Dcp-1 antibody (7 days after egg laying). (E) Quantification of dying cells in GFP positive clone area in (A-D). Error bars, SD; ***, p<0.001, ****, p<0.0001 by one-way ANOVA multiple comparison test. (F-H) Eye-antennal disc bearing GFP-labeled clones of indicated genotypes (7 days after egg laying). (I) Quantification of clone size (% of total clone area per disc area in eye-antennal disc) of indicated genotype. Error bars, SD; n.s., p>0.05 (not significant), *, p<0.05, p<0.0001 by two-tailed student’s t test. (J-M) Eye-antennal disc bearing GFP-labeled clones of indicated genotypes stained with anti-cleaved Dcp-1 antibody (5 days after egg laying). (N) Quantification of dying cells in GFP positive clone area in (J-M). Error bars, SD; n.s., p>0.05 (not significant) by one-way ANOVA multiple comparison test.

### miR-306 and miR-79 suppress tumor growth by enhancing JNK signaling

We thus examined whether Ras activation or cell polarity defect cooperates with miR-306 or miR-79 to induce cell death. Overexpression of the miR-9c/306/79/9b cluster, miR-306, or miR-79 in Ras^V12^-expresing clones did not affect their growth (Figure 3—figure supplement 1A-D, quantified in Figure 3—figure supplement 1E), indicating that Ras signaling does not cooperate with these miRNAs. Notably, overexpression of these miRNAs in *dlg^−/−^* clones significantly reduced their clone size (Figure 3A-D, quantified in Figure 3I). In addition, blocking cell death by overexpression of p35 cancelled the ability of these miRNAs to reduce *dlg^−/−^* clone size (Figure 3E-H, quantified in Figure 3I), suggesting that these miRNAs block *dlg^−/−^* clone growth by promoting cell death. These data show that miR-306 or miR-79 cooperates with loss of cell polarity to induce synthetic lethality.

**Figure 3.**
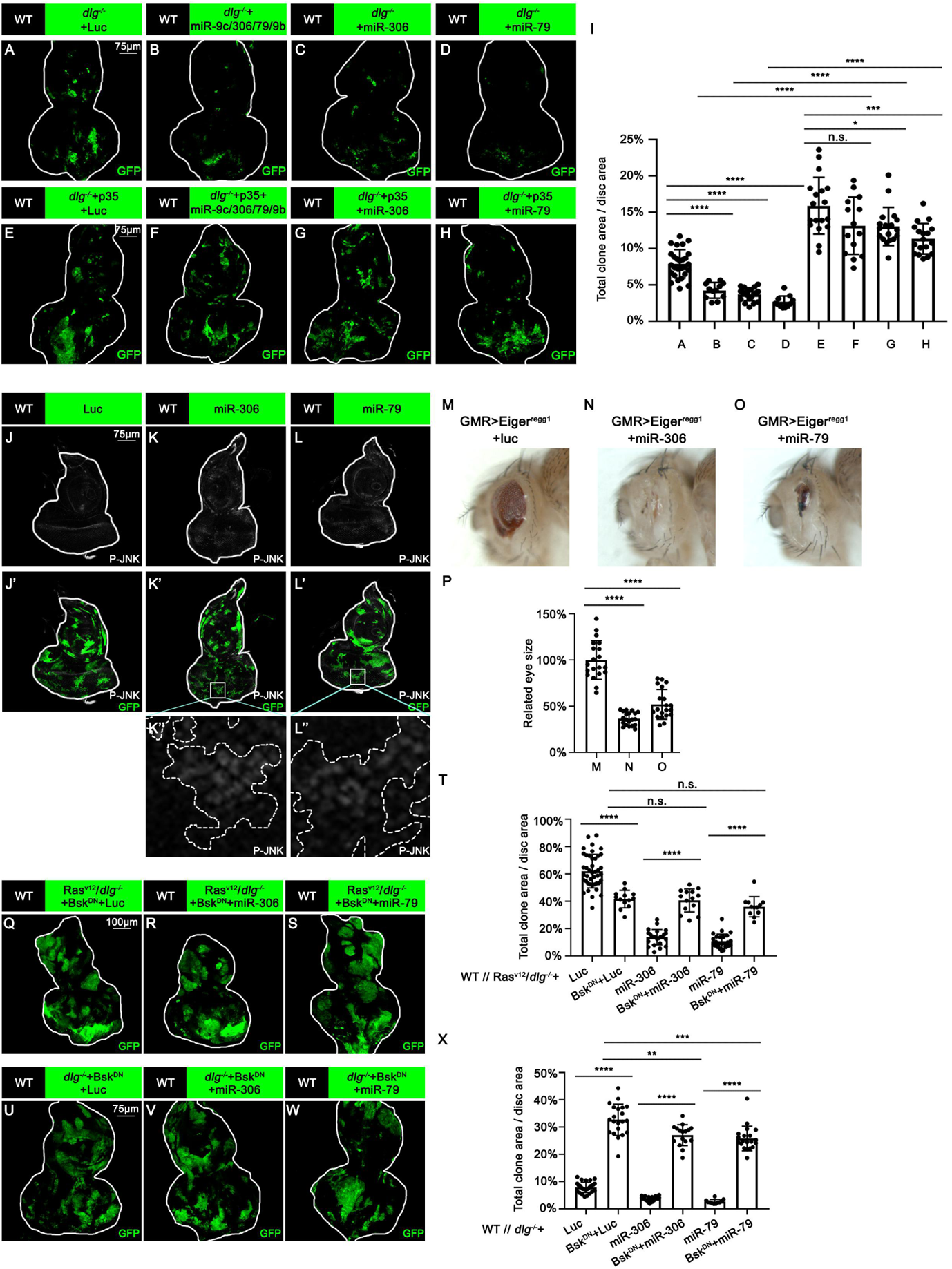
miR-306 and mir-79 suppress tumor growth and promote cell competition by promoting JNK signaling. (A-H) Eye-antennal disc bearing GFP-labeled clones of indicated genotypes (5 days after egg laying). (I) Quantification of clone size (% of total clone area per disc area in eye-antennal disc) of (A-H). Error bars, SD; n.s., p>0.05 (not significant), *, p<0.05, ***, p<0.001, ****, p<0.0001 by two-tailed student’s t test. (J-L) Eye-antennal disc bearing GFP-labeled clones of indicated genotypes stained with anti-Phospho-JNK antibody (5 days after egg laying). (M-O) Adult eye phenotype of flies with indicated genotypes. (P) Quantification of adult eye size (normalized to control) of (M-O). Error bars, SD; ****, p<0.0001 by one-way ANOVA multiple comparison test. (Q-S) Eye-antennal disc bearing GFP-labeled clones of indicated genotypes (6 days after egg laying). (T) Quantification of clone size (% of total clone area per disc area in eye-antennal disc) of indicated genotype. Error bars, SD; n.s., p>0.05 (not significant), ****, p<0.0001 by two-tailed student’s t test. (U-W) Eye-antennal disc bearing GFP-labeled clones of indicated genotypes (5 days after egg laying). (X) Quantification of clone size (% of total clone area per disc area in eye-antennal disc) of indicated genotype. Error bars, SD; **, p<0.01, **, p<0.001, ****, p<0.0001 by two-tailed student’s t test.

We then sought to identify the polarity defect-induced intracellular signaling that cooperates with miR-306 or miR-79 to induce cell death. It has been shown that clones of cells mutant for cell polarity genes such as *dlg* activate JNK signaling via the *Drosophila* tumor necrosis factor (TNF) Eiger (Brumby & Richardson, 2003; Igaki et al., 2009). We found that overexpression of miR-306 or miR-79 alone moderately activated JNK signaling in the eye-antennal discs, as visualized by anti-p-JNK antibody staining and the *puc-LacZ* reporter (Figure 3J-L, Figure 3—figure supplement 2A-C). In addition, Western blot analysis with anti-p-JNK antibody revealed that overexpression of miR-306 or miR-79 in the eyes using the GMR-Gal4 driver caused JNK activation (Figure 3—figure supplement 2D). Notably, although overexpression of miR-306 or miR-79 alone in the eyes had no significant effect on eye morphology (Figure 3—figure supplement 2E-G, quantified in Figure 3—figure supplement 2H), they dramatically enhanced the reduced-eye phenotype caused by overexpression of Eiger (Figure 3M-O, quantified in Figure 3P). It has been shown that the severity of the reduced-eye phenotype depends on the levels of JNK activation and subsequent cell death (Igaki et al., 2002; Igaki et al., 2006; Igaki et al., 2009), suggesting that miR-306 and miR-79 enhance Eiger-mediated activation of JNK signaling. Indeed, blocking JNK signaling by overexpression of a dominant-negative form of *Drosophila* JNK Basket (Bsk^DN^) cancelled the tumor-suppressive activity of miR-306 or miR-79 against Ras^V12^/*dlg^−/−^* or Ras^V12^/*lgl^−/−^* tumors (Figure 3Q-S, quantified in Figure 3T and Figure 3—figure supplement 3A-C, quantified in Figure 3—figure supplement 3D). Moreover, overexpression of Bsk^DN^ significantly increased the size of *dlg^−/−^* or *lgl^−/−^* clones overexpressing miR-306 or miR-79 (Figure 3U-W, quantified in Figure 3X; Figure 3—figure supplement 3E-J, quantified in Figure 3—figure supplement 3K). Together, these data suggest that miR-306 and miR-79 suppress growth of malignant tumors by enhancing JNK signaling activation.

### miR-306 and miR-79 enhance JNK signaling in different types of tumors

We next examined whether miR-306 or miR-79 suppresses growth of other types of tumors with elevated JNK signaling via an Eiger-independent mechanism. Overexpression of an activated form of the *Drosophila* PDGF/VEGF receptor homolog (PVR^act^) results in JNK activation and tumor formation in the wing disc (C. W. Wang, Purkayastha, Jones, Thaker, & Banerjee, 2016) and eye-antennal disc (Figure 4A). This tumor growth was significantly suppressed by overexpression of the miR-9c/306/79/9b cluster, miR-306, or miR-79 (Figure 4B-D, quantified in Figure 4E). In addition, the size of clones overexpressing the oncogene Src64B in the eye-antennal disc (Figure 4F), which activate JNK signaling (Enomoto & Igaki, 2013), was significantly reduced when the miR-9c/306/79/9b cluster, miR-306, or miR-79 was coexpressed (Figure 4G-I, quantified in Figure 4J). Moreover, non-autonomous overgrowth of surrounding wild-type tissue by Src64B-overexpressing clones (Enomoto & Igaki, 2013) was significantly suppressed by coexpression of these miRNAs (Figure 4K-N, quantified in Figure 4O). Furthermore, the size of clones mutant for an RNA helicase Hel25E or an adaptor protein Mahj, both of which are eliminated by JNK-dependent cell death when surrounded by wild-type cells (Nagata, Nakamura, Sanaki, & Igaki, 2019; Tamori et al., 2010), was significantly reduced when these miRNAs were coexpressed (Figure 4—figure supplement 1A-D, quantified in Figure 4—figure supplement 1E and Figure 4—figure supplement 1 F-I, quantified in Figure 4—figure supplement 1J). These data suggest that miR-306 and miR-79 broadly enhance JNK signaling activity stimulated by different upstream signaling.

**Figure 4.**
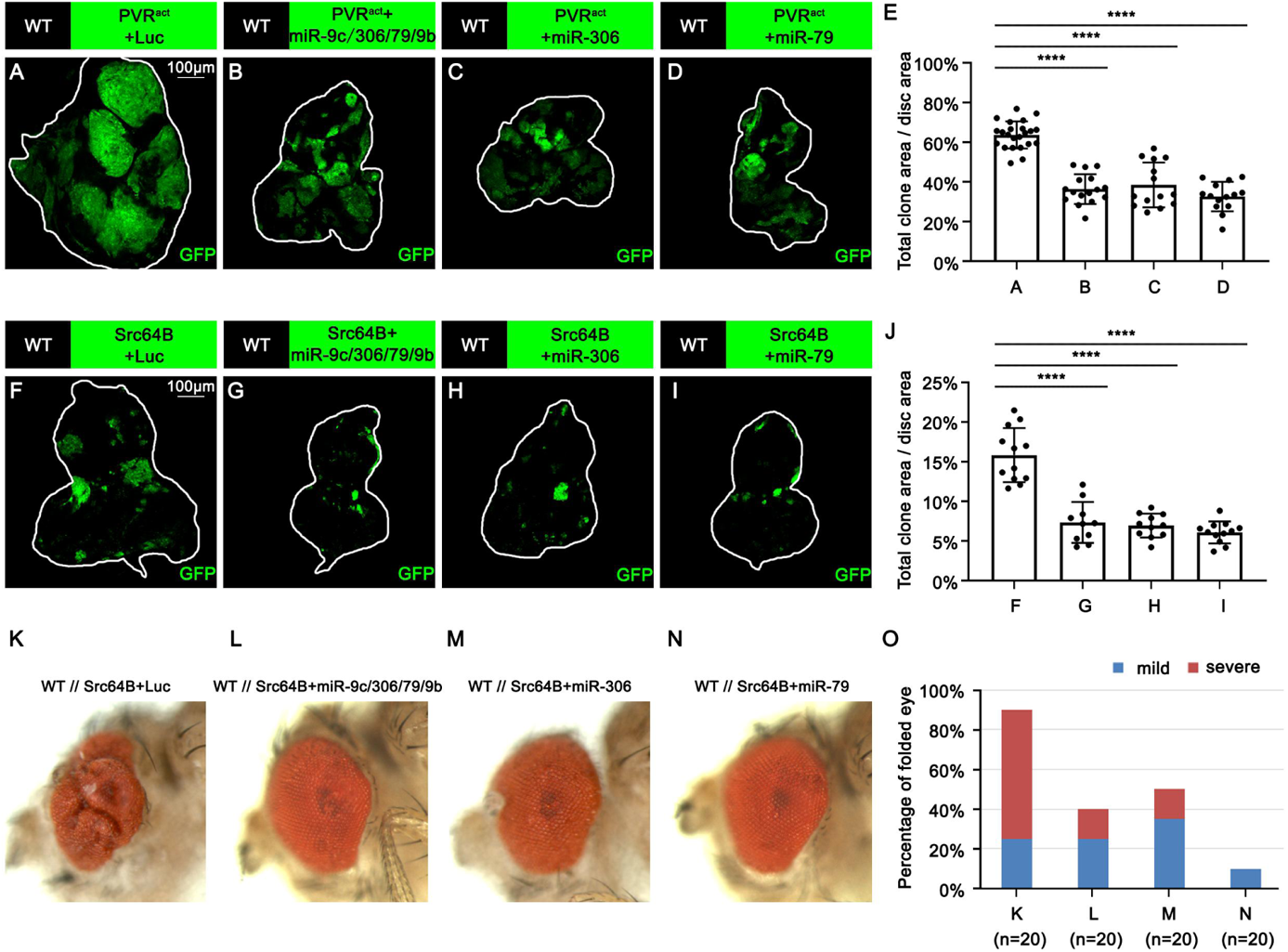
miR-306 and miR-79 suppress growth of multiple types of tumor models. (A-D) Eye-antennal disc bearing GFP-labeled clones of indicated genotypes (7 days after egg laying). (E) Quantification of clone size (% of total clone area per disc area in eye-antennal disc) of (A-D). Error bars, SD; ****, p<0.0001 by one-way ANOVA multiple comparison test. (F-I) Eye-antennal disc bearing GFP-labeled clones of indicated genotypes (6 days after egg laying). (J) Quantification of clone size (% of total clone area per disc area in eye-antennal disc) of (F-I). Error bars, SD; ****, p<0.0001 by one-way ANOVA multiple comparison test. (K-N) Adult eye phenotype of flies with indicated genotypes. (O) Quantification of percentage of folded eye in (K-N). n=20 for each group.

### miR-306 and miR-79 enhance JNK signaling activity by targeting dRNF146

We next sought to identify the mechanism by which miR-306 and miR-79 enhance JNK signaling by searching for the target gene(s) of these miRNAs. The clustered miRNAs often target overlapping sets of genes and thus co-regulate various biological processes (Kim et al., 2009; Y. Wang, Luo, Zhang, & Lu, 2016; Yuan et al., 2009). Given that miR-306 and miR-79 are located on the same miRNA cluster, we searched for the common targets of these miRNAs using the online software TargeyScanFly 7.2 (http://www.targetscan.org/fly_72/) and found 11 mRNAs that were predicted to be targets of both miR-306 and miR-79 (Figure 5A). We then examined whether knocking down of each one of these candidate genes could activate JNK signaling in *Drosophila* wing discs, where a clear JNK activation was observed when miR-306 or miR-79 was overexpressed (Figure 5—figure supplement 1). As a result, we found that knocking down of dRNF146, but not any other available RNAis for the candidate genes, resulted in JNK activation (Figure 5B-C, Figure 5—figure supplement 2). The dRNF146 mRNA had putative target sites of miR-306 and miR-79 in its 3’UTR region (Figure 5D). To confirm that dRNF146 mRNA is a direct target of miR-306 and miR-79, we performed a dual-luciferase reporter assay in *Drosophila* S2 cells using wild-type dRNF146 3’UTR (dRNF146 WT) or mutant dRNF146 3’UTR bearing mutations at the putative binding site of miR-306 (dRNF146 m1) or miR-79 (dRNF146 m2) (Figure 5D). We found that miR-306 and miR-79 reduced wild-type dRNF146 3’UTR expression but did not affect respective mutant dRNF146 3’UTR (Figure 5E), indicating that miR-306 and miR-79 directly target dRNF146 3’UTR (Figure 5D). We also confirmed that overexpression of miR-306 or miR-79 reduced the endogenous levels of dRNF146 protein in *Drosophila* S2 cells (Figure 5F).

**Figure 5.**
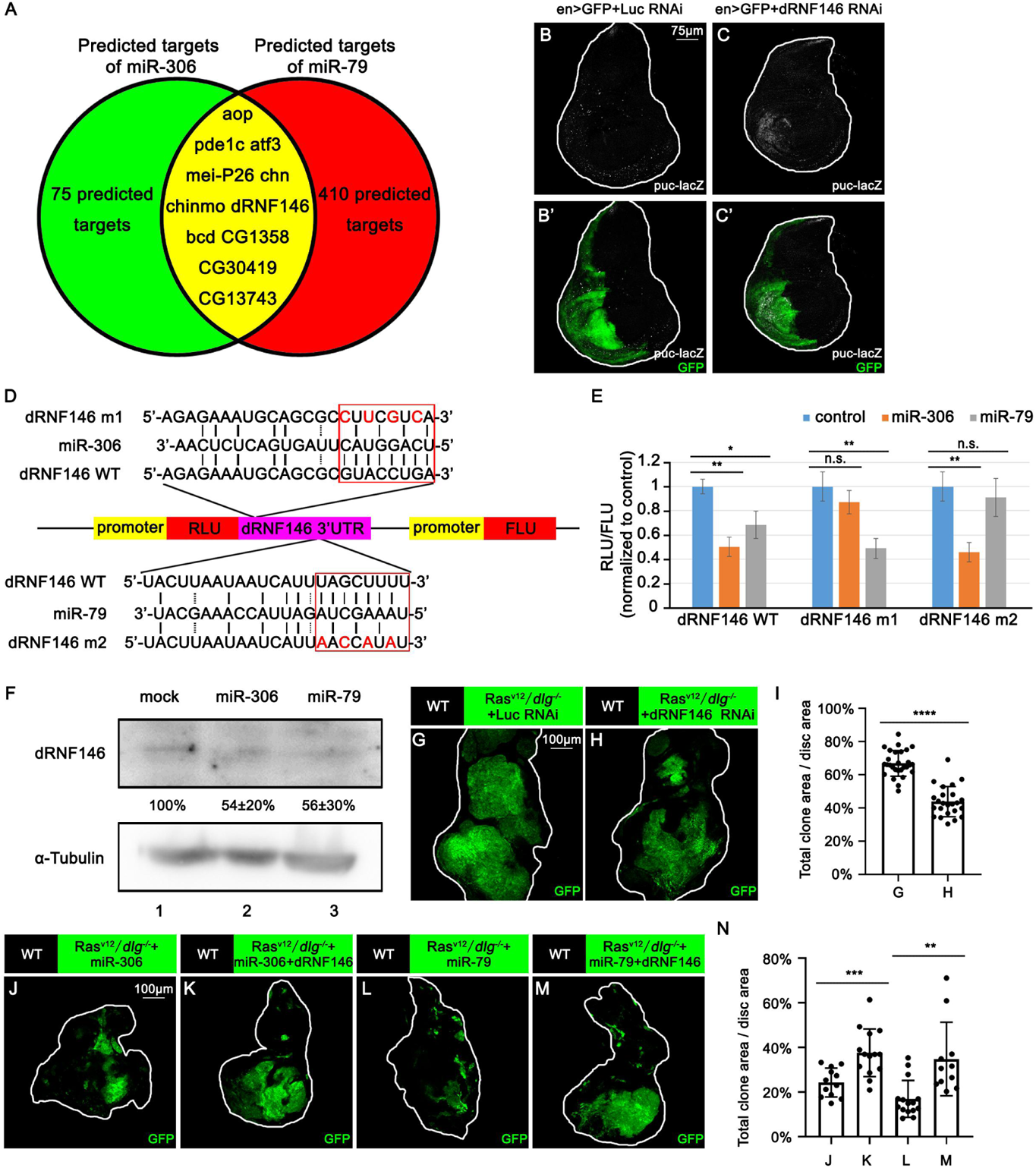
miR-306 and mir-79 suppress tumor growth and promote cell competition by targeting dRNF146. (A) Predicted targets of miR-306 and miR-79. (B-C) Wing disc of indicated genotypes with puc-lacZ background stained with anti-β-galactosidase antibody (5 or 6 days after egg laying). (D) Schematic of the wild-type and mutation-type 3’UTR vector with miRNA binding sites for miR-306 and miR-79, respectively. Red letters shows the mutation sites. Red box shows the seed sequence pairing region. (E) RLU/FLU rate from dual luciferase assay. n=3, error bars, SD; n.s., p>0.05 (not significant), **, p<0.01 by two-tailed student’s t test. (F) *Drsophila* S2 cells were transfected with empty plasmid or plasmid expressing miR-306 or miR-79. After 48 hours, cell lysates were subjected to Western blots using indicated antibodies. The relative levels of dRNF146 protein were presented as means ± SD (normalized to lane 1) from three independent experiments. (G-H) Eye-antennal disc bearing GFP-labeled clones of indicated genotypes (7 days after egg laying). (I) Quantification of clone size (% of total clone area per disc area in eye-antennal disc) of (G-H). Error bars, SD; ****, p<0.0001 by two-tailed student’s t test. (J-M) Eye-antennal disc bearing GFP-labeled clones of indicated genotypes (7 days after egg laying). (N) Quantification of clone size (% of total clone area per disc area in eye-antennal disc) of (J-M). Error bars, SD; **, p<0.01, ***, p<0.001 by two-tailed student’s t test.

We next investigated whether dRNF146 is the responsible target of miR-306 and miR-79 for the enhancement of JNK signaling. We found that, while knockdown of dRNF146 did not affect normal tissue growth (Figure 5—figure supplement 3 A-B, quantified in Figure 5—figure supplement 3C), it significantly suppressed Ras^V12^/*dlg^−/−^* tumor growth (Figure 5G-H, quantified in Figure 5I) and promoted elimination of *dlg^−/−^* clones (Figure 5—figure supplement 3D-E, quantified in Figure 5—figure supplement 3F). In addition, overexpression of dRNF146 rescued the reduced-eye phenotype caused by overexpression of miR-306 or miR-79 in the eyes (Figure 5—figure supplement 3G-J, quantified in Figure 5—figure supplement 3K). Moreover, knocking down of dRNF146 significantly enhanced Eiger-induced reduced-eye phenotype (Figure 5—figure supplement 3L-O, quantified in Figure 5—figure supplement 3P). Furthermore, overexpression of dRNF146 cancelled the tumor-suppressive effect of miR-306 or miR-79 on Ras^V12^/*dlg^−/−^* tumors (Figure 5J-M, quantified in Figure 5N). The dRNF146 overexpression also cancelled the enhanced elimination of *dlg^−/−^* clones by miR-306 or miR-79 (Figure 5—figure supplement 3Q-T, quantified in Figure 5—figure supplement 3U). Together, these data indicate that miR-306 and miR-79 directly target dRNF146 mRNA, thereby enhancing JNK signaling activity and thus exerting the tumor-suppressive effects.

### DRNF146 promotes Tnks degradation

We next investigated the mechanism by which downregulation of dRNF146 by miR-306 or miR-79 enhances JNK signaling activity. It has been shown in *Drosophila* embryos, larvae, wing discs, and adult eyes that loss of dRNF146 upregulates the protein levels of Tnks (Gultekin & Steller, 2019; Z. Wang et al., 2019), a poly-ADP-ribose polymerase that mediates K63-linked poly-ubiquitination of JNK and thereby promotes JNK-dependent apoptosis in *Drosophila* (Feng et al., 2018; Li et al., 2018). In addition, loss of dRNF146 was shown to enhance rough-eye phenotype caused by Tnks overexpression (Gultekin & Steller, 2019). These observations raise the possibility that downregulation of dRNF146 by miR-306 or miR-79 enhances JNK signaling via upregulation of Tnks. Indeed, as reported previously (Feng et al., 2018; Li et al., 2018), Western blot analysis revealed that overexpression of Tnks induces phosphorylation of JNK (JNK activation) in S2 cells (Figure 6A, lane 2 vs. lane 1). Notably, coexpression of dRNF146 significantly downregulated Tnks protein level and suppressed Tnks-induced JNK phosphorylation (Figure 6A, lane 3 vs. lane 2). Moreover, knocking down of dRNF146 significantly upregulated Tnks protein level and promoted JNK phosphorylation (Figure 6B).

**Figure 6.**
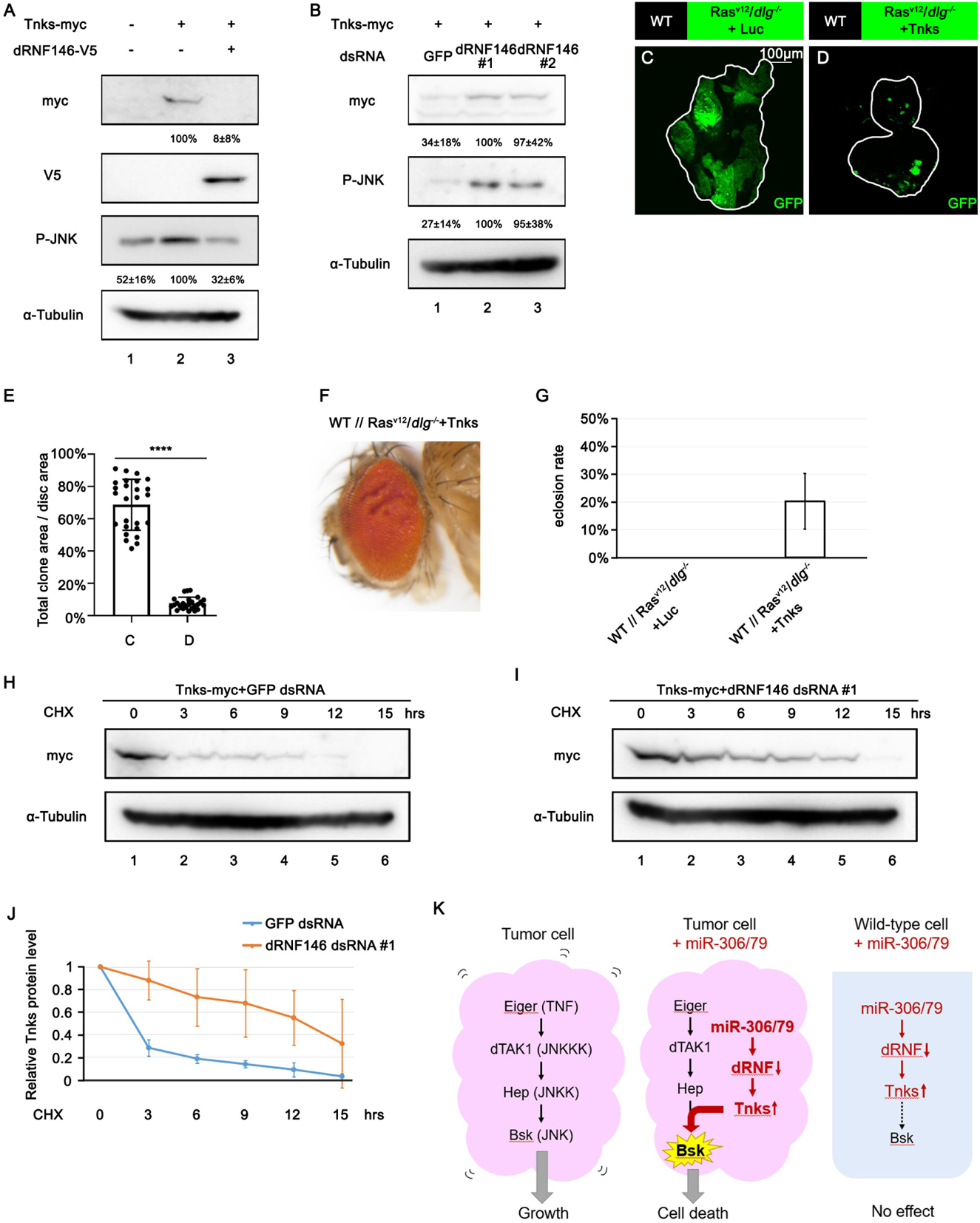
DRNF146 promotes poly-ubiquitination and degradation of Tnks. (A) *Drsophila* S2 cells were transfected with plasmids expressing indicated proteins. Cell lysates were subjected to Western blots using indicated antibodies. The relative levels of myc-tagged Tnks or P-JNK protein were presented as means ± SD (normalized to lane 2) from three independent experiments. (B) *Drsophila* S2 cells were transfected with plasmid expressing indicated protein and dsRNA targeting indicated gene. Cell lysates were subjected to Western blots using indicated antibodies. The relative levels of myc-tagged Tnks or P-JNK protein were presented as means ± SD (normalized to lane 2) from three independent experiments. (C-D) Eye-antennal disc bearing GFP-labeled clones of indicated genotypes (7 days after egg laying). (E) Quantification of clone size (% of total clone area per disc area in eye-antennal disc) of (A-B). Error bars, SD; ****, p<0.0001 by two-tailed student’s t test. (F) Adult eye phenotype of flies with indicated genotypes. (G) Eclosion rate of flies with indicated genotypes. Data from three independent experiment, n>30 for each group in one experiment; error bars, SD. (H-I) *Drsophila* S2 cells were transfected with plasmid expressing indicated protein and dsRNA targeting indicated gene. After 36 hours, cells were treated with 50 μg/ml CHX for the indicated periods. Cell lysates were subjected to Western blots using indicated antibodies. (J) The relative levels of Tnks protein shown in (H) and (I) were plotted. All data represent means and SD of three independent experiments. (K) A model for tumor elimination by miR-306/79. Tumor cell with elevated canonical JNK signaling via Eiger/TNF, dTAK1/JNKKK and Hep/JNKK grows in a Bsk/JNK-dependent manner. Overexpression of miR-306 or miR-79 in JNK-activated tumor cell results in overactivation of JNK signaling to the lethal level via dRNF-Tnks-mediated non-canonical JNK-activating signaling. Overexpression of miR-306 or miR-79 in normal cells has no significant effect on JNK signaling.

These data support the notion that downregulation of dRNF146 enhances JNK activation via upregulation of Tnks. Indeed, overexpression of Tnks was sufficient to suppress growth of Ras^V12^/*dlg^−/−^* tumors (Figure 6C-D, quantified in Figure 6E) and rescue the lethality of flies bearing Ras^V12^/*dlg^−/−^* tumors in the eye-antennal discs (Figure 6F-G).

Finally, we sought to clarify the mechanism by which downregulation of dRNF146 upregulates Tnks. The upregulation of Tnks can be caused by either elevated Tnks protein synthesis or reduced Tnks protein degradation. We thus examined the possibility that dRNF146 promotes degradation of Tnks. Blocking new protein synthesis in S2 cells by the protein synthesis inhibitor cycloheximide (CHX) resulted in a time-dependent depletion of Tnks protein with a half-life of less than 3 hours (Figure 6H, quantified in Figure 6J). This depletion of Tnks was significantly retarded when dRNF146 was knocked down (Figure 6I, quantified in Figure 6J). These data indicate that endogenous dRNF146 promotes degradation of Tnks protein. Taken together, our data show that miR-306 or miR-79 directly targets dRNF146, thereby leading to elevation of Tnks protein that induces non-canonical activation of JNK signaling (Figure 6K).

## Discussion

In this study, we have identified the clustered miRNAs miR-306 and miR-79 as novel anti-tumor miRNAs that selectively eliminate JNK-activated tumors from *Drosophila* imaginal epithelia. Mechanistically, miR-306 and miR-79 directly target dRNF146, an E3 ligase that promotes degradation of a poly-ADP-ribose polymerase Tnks, thereby leading to upregulation of Tnks and thus promoting JNK activation (Figure 6K). Importantly, this non-canonical mode of JNK activation has only a weak effect on normal tissue growth but it strongly blocks tumor growth by overactivating JNK signaling when tumors already possess elevated JNK signaling via the canonical JNK pathway (Figure 6K). Given that tumors or pre-malignant mutant cells often activate canonical JNK signaling, miR-306 and miR-79 can be novel ideal targets of cancer therapy.

Our study identified several putative co-target genes of miR-306 and miR-79 (Figure 5A). Interestingly, some of these genes (*Atf3*, *chinmo*, and *chn*) have been reported to be involved in tumor growth in *Drosophila*. *Atf3* encodes an AP-1 transcription factor which was shown to be a polarity-loss responsive gene acting downstream of the membrane-associated Scrib polarity complex (Donohoe et al., 2018). Knockdown of *Atf3* suppresses growth and invasion of Ras^V12^/*scrib^−/−^* tumors in eye-antennal discs (Atkins et al., 2016). *Chinmo* is a BTB-zinc finger oncogene that is up-regulated by JNK signaling in tumors (Doggett et al., 2015). Although loss of *chinmo* does not significantly suppress tumor growth, overexpression of *chinmo* with Ras^V12^ or an activated Notch is sufficient to promote tumor growth in eye-antennal discs (Doggett et al., 2015). *Chn* encodes a zinc finger transcription factor that cooperates with *scrib^−/−^* to promote tumor growth (Turkel et al., 2013). Although we found that knockdown of these genes did not activate JNK signaling, it is possible that these putative target genes also contribute to the miR-306/miR-79-induced tumor suppression.

Intriguingly, it has been reported that miR-79 is down-regulated in Ras^V12^/*lgl-RNAi* tumors in *Drosophila* wing discs (Shu et al., 2017). Given that miR-306 is located in the same miRNA cluster with miR-79, it is highly possible that miR-306 is also down-regulated in tumors. This suggests that tumors have the mechanism that downregulates anti-tumor miRNAs for their survival and growth. Future studies on the mechanism of how tumors regulate these miRNAs would provide new understanding of tumor biology.

Our study uncovered the miR-306/79-dRNF146-Tnks axis as non-canonical JNK enhancer that selectively eliminates JNK-activated tumors in *Drosophila*. Considering that miR-9, the mammalian homolog of miR-79, is predicted to target mammalian RNF146 (Figure 5—figure supplement 4) and that JNK signaling is highly conserved throughout evolution, it opens up the possibility of developing a new miRNA-based strategy against cancer.

## Materials and Methods

### Fly stocks

All flies used were reared at 25°C on a standard cornmeal/yeast diet. Fluorescently labelled mitotic clones were produced in larval imaginal discs using the following strains: Tub-Gal80, FRT40A; eyFLP6, Act>y^+^>Gal4, UAS-GFP (40A tester), FRT42D, Tub-Gal80/CyO; eyFLP6, Act>y^+^>Gal4, UAS-GFP (42D tester), Tub-Gal80, FRT19A; eyFLP5, Act>y^+^>Gal4, UAS-GFP (19A tester #1), Tub-Gal80, FRT19A; eyFLP6, Act>y^+^>Gal4, UAS-GFP (19A tester #2). Additional strains used are the following: dlg^m52^ (Goode & Perrimon, 1997), puc-lacZ (Igaki et al., 2006), UAS-Ras^v12^ (Igaki et al., 2006), UAS-Bsk^DN^ (Adachi-Yamada et al., 1999), UAS-Src64B (Wills, Bateman, Korey, Comer, & Van Vactor, 1999), Hel25E^ccp-8^ (Nagata et al., 2019), Mahj^1^ (Tamori et al., 2010), UAS-N^act^ (Hori et al., 2004), UAS-dRNF146 (Gultekin & Steller, 2019); lgl^4^ (BDSC #36289), UAS-p35 (BDSC #5073), UAS-PVR^act^ (BDSC #58496), UAS-Yki^S168A^ (BDSC #28836), UAS-Luciferase (BDSC #35788), UAS-bantam (BDSC #60672), UAS-miR-9c,306,79,9b (BDSC #41156), UAS-miR-79 (BDSC #41145), UAS-miR-2a-2,2a-1,2b-2 (BDSC #59849), UAS-miR-2b-1 (BDSC #41128), UAS-miR-7 (BDSC #41137), UAS-miR-8 (BDSC #41176), UAS-miR-9a (BDSC #41138), UAS-miR-9b (BDSC #41131), UAS-miR-9c (BDSC #41139), UAS-miR-11 (BDSC #59865), UAS-miR-12 (BDSC #41140), UAS-miR-13a,13b-1,2c (BDSC #64097), UAS-miR-13b-2 (BDSC #59867), UAS-miR-14 (BDSC #41178), UAS-miR-34 (BDSC #41158), UAS-miR-92a (BDSC #41153), UAS-miR-124 (BDSC #41126), UAS-miR-184 (BDSC #41174), UAS-miR-252 (BDSC #41127), UAS-miR-276a (BDSC #41143), UAS-miR-276b (BDSC #41162), UAS-miR-278 (BDSC #41180), UAS-miR-279 (BDSC #41147), UAS-miR-282 (BDSC #41165), UAS-miR-305 (BDSC #41152), UAS-miR-310 (BDSC #41155), UAS-miR-317 (BDSC #59913), UAS-miR-958 (BDSC #41222), UAS-miR-975,976,977 (BDSC #60635), UAS-miR-981 (BDSC #60639), UAS-miR-984 (BDSC #41224), UAS-miR-988 (BDSC #41196), UAS-miR-995 (BDSC #41199), UAS-miR-996 (BDSC #60653), UAS-miR-998 (BDSC #63043), UAS-Luciferase RNAi (BDSC #31603), UAS-aop RNAi (BDSC #34909), UAS-pde1c RNAi (BDSC #55925), UAS-atf3 RNAi (BDSC #26741), UAS-mei-P26 RNAi (BDSC #57268), UAS-chn RNAi (BDSC #26779), UAS-chinmo RNAi (BDSC #26777), UAS-dRNF146 RNAi (BDSC #40882), UAS-bcd RNAi (BDSC #33886) and UAS-CG1358 RNAi (BDSC #64848) from Bloomington *Drosophila* Stock Center; UAS-miR-306 (FlyORF #F002214) from FlyORF; UAS-Tnks from Core Facility of Drosophila Resource and Technology, Center for Excellence in Molecular Cell Science, Chinese Academy of Sciences.

### Clone size measurement

Eye-antennal disc images were taken with a Leica SP8 confocal microscope or Olympus Fluoview FV3000 confocal microscope. To measure clone size, ImageJ (Fiji) software was used to determine the threshold of the fluorescence. Total clone area/disc area (%) in the eye-antennal disc was calculated using ImageJ and Prism 8 (Graphpad).

### Histology

Larval tissues were stained with standard immunohistochemical procedures using rabbit anti-Phospho-JNK polyclonal antibody (Cell Signaling Technology, Cat #4668, 1:100), chicken anti-β-galactosidase antibody (Abcam, Cat #ab9361, 1:1000), rabbit anti-Cleaved Drosophila Dcp-1 (Asp216) antibody (Cell Signaling Technology, Cat#9578, 1:100), goat anti-rabbit secondary antibody, Alexa Fluor 647 (Thermo Fisher Scientific, Cat #A32733, 1:250) or goat anti-chicken secondary antibody, Alexa Fluor 647 (Thermo Fisher Scientific, Cat #A21449, 1:250). Samples were mounted with DAPI-containing SlowFade Gold Antifade Reagent (Thermo Fisher Scientific, Cat #S36937). Images were taken with a Leica SP8 confocal microscope.

### Plasmid and *in vitro* transcription of dsRNA

pAc5.1/V5-His vector (Thermo Fisher Scientific, Cat #V411020) was used to construct plasmids for expressing proteins or miRNAs in *Drosophila* S2 cells. The dRNF146 or Tnks ORF was amplified from fly cDNAs via PCR. The dRNF146 ORF was cloned into the EcoR І-Xho І site of the pAc5.1/V5-His vector. The Tnks ORF carrying a myc tag at its 5’-end was cloned into the Kpn І-Xho І site of the pAc5.1/V5-His vector. Extended region of miR-306 (−184 to +136) or miR-79 (−124 to +131) was amplified from fly cDNAs via PCR and cloned into the Kpn І-EcoR І site of the pAc5.1/V5-His vector.

dRNF146 dsRNA #1 and #2, respectively targeting the 1-318 and 319-667 region of dRNF146 ORF, used for dRNF146 RNAi were transcribed *in vitro* using T7 RNA polymerase (Promega, Cat #P2075) at 37°C for 4 hrs from the PCR products.

### Cell culture and transfection

*Drosophila* S2 cells were grown and maintained in Schneider’s *Drosophila* medium (Thermo Fisher Scientific, Cat #21720024)/10% fetal bovine serum (FBS) and penicillin/streptomycin. *Drosophila* S2 cells were plated in 100-mm plates or six-well plates and grown overnight to reach 70% confluence. After that, DNA plasmids or dsRNAs were transfected into the cells using FuGene HD transfection reagent (Promega, Cat #PRE2311) according to the manufacturer’s protocol.

### Inhibitors

The protein synthesis inhibitor CHX (Santa Cruz Biotechnology, Cat #SC-3508) was used at 50 μg/ml. The proteasome inhibitor MG-132 (Sigma Aldrich, Cat #C2211) was used at 50 μM. The proteasome inhibitor lactacystin (Peptide Institute, Code #4369-v) was used at 10μM. The vacuolar H^+^-ATPase inhibitor Bafilomycin A1 (Bioviotica, Cat #BVT-0252) was used at 100nM. The amphisome-lysosome fusion inhibitor leupeptin (MedChemExpress, Cat #HY-18234A) was used at 50μM.

### Western blots

Cultured Drosophila S2 cells were harvested and then lysed in cell lysis buffer. The cell lysates were then subjected to SDS-PAGE, followed by Western blots using anti-α-Tubulin monoclonal antibody (Sigma Aldrich, Cat #T5168, 1:5000), anti-Phospho-JNK polyclonal antibody (Cell Signaling Technology, Cat #9251, 1:1000), anti-dRNF146 polyclonal antibody (gift from Prof. Hermann Steller, 1:100) (Gultekin & Steller, 2019), anti-V5 tag monoclonal antibody (Thermo Fisher Scientific, Cat #R960-25, 1:5000), anti-myc tag polyclonal antibody (MBL, Code #562, 1:1000), anti-mouse IgG, HRP-linked antibody (Cell Signaling Technology, Cat #7076,1:5000), anti-rabbit IgG, HRP-linked antibody (Cell Signaling Technology, Cat #7074, 1:5000) or anti-guinea pig IgG, HRP-conjugate antibody (Thermo Fisher Scientific, Cat #A18769, 1:1000).

### Dual-luciferase reporter assay

The psiCHECK-2 vector (Promega, Cat #C8021) was used to construct plasmids for dual-luciferase reporter assay. dRNF146 3’UTR or its mutant was cloned into the Xho І-Not I site of the psiCHECK-2 vector. Renilla luciferase activity and firefly luciferase activity were measured using GloMax-Multi Jr Single-Tube Multimode Reader (Promega) according to the manufacturer’s protocol.

### Statistical analysis

When comparing two groups, statistical significance was tested using a student’s t-test. When comparing multiple groups, statistical significance was tested using a one-way ANOVA multiple comparison test. In all figures, significance is indicated as follows: p>0.05 is indicated by n.s. (not significant), p < 0.05 is indicated by *, p < 0.01 is indicated by **, p < 0.001 is indicated by ***, and p<0.0001 is indicated by ****.

## Acknowledgements

We thank M. Matsuoka and K. Gomi for technical support, Hermann Steller, the Bloomington Drosophila Stock Center, the National Institute of Genetics Stock Center (NIG-FLY), the Drosophila Genomics and Genetic Resources (DGGR, Kyoto Institute of Technology), the Vienna Drosophila Resource Center (VDRC), and the Core Facility of Drosophila Resource and Technology, Center for Excellence in Molecular Cell Science, Chinese Academy of Sciences for fly stocks and reagents. We also thank members of the Igaki laboratory for discussions. This work was supported by grants from the MEXT/JSPS KAKENHI (Grant Number 20H05320, 21H05284, and 21H05039) to T.I, Japan Agency for Medical Research and Development (Project for Elucidating and Controlling Mechanisms of Aging and Longevity; Grant Number 20gm5010001) to T.I, the Takeda Science Foundation to T.I, and the Naito Foundation to T.I. Z.W was supported by JSPS Postdoctoral Fellowships for Research in Japan, X.X was supported by China Scholarship Council for Research in Japan.

## Figures

**Figure 1—figure supplement 1.**
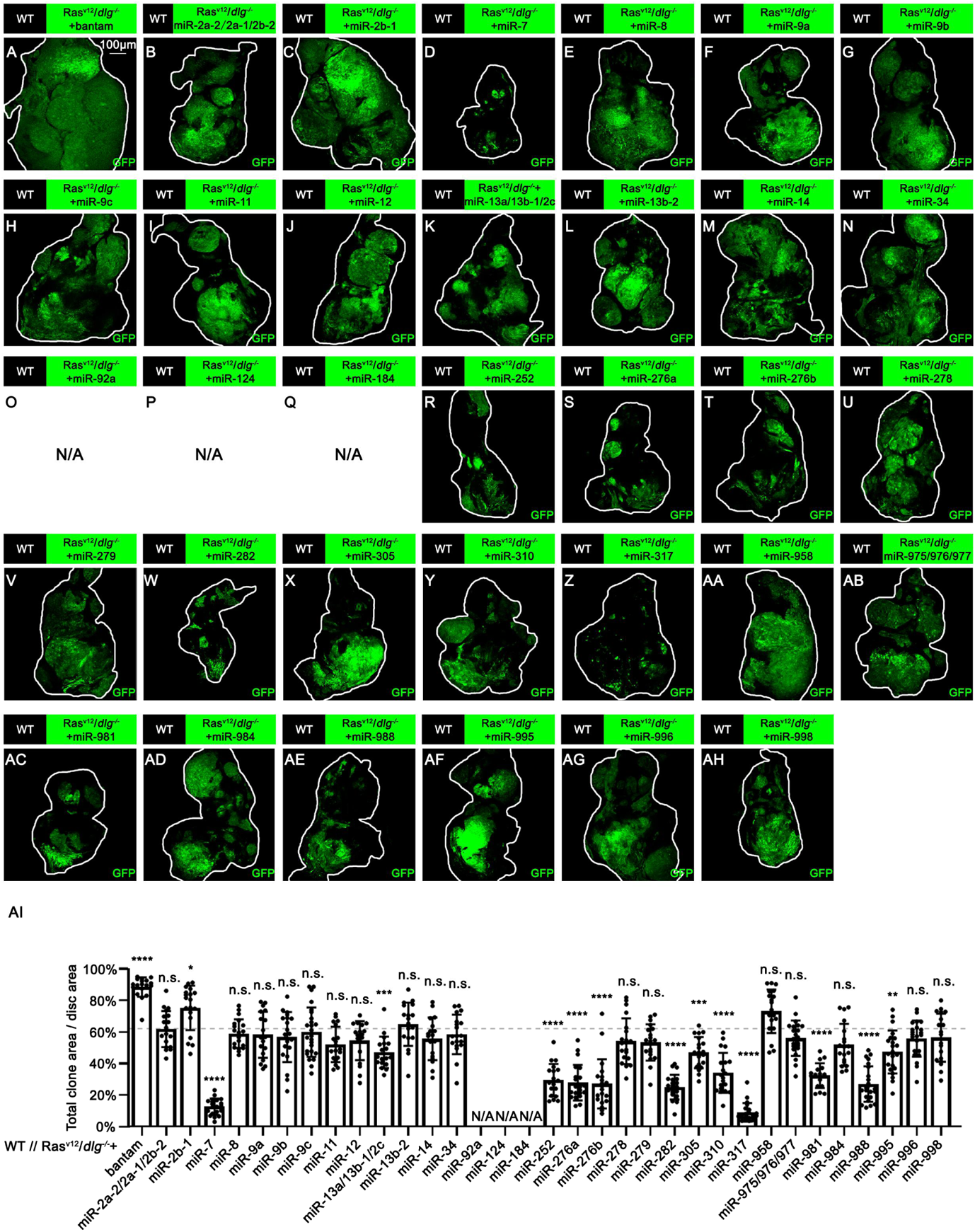
Effect of miRNAs or miRNA clusters on Ras^V12^/*dlg^−/−^* tumor growth. (A-AH) Eye-antennal disc bearing GFP-labeled clones of indicated genotypes (7 days after egg laying). (AI) Quantification of clone size (% of total clone area per disc area in eye-antennal disc) of (A-AH). Dotted line shows the relative clone size of WT//Dlg^−/−^+Ras^v12^+Luc in average (Control). Error bars, SD; n.s., p>0.05 (not significant), *, p<0.05, **, p<0.01, ***, p<0.001, ****, p<0.0001 by one-way ANOVA multiple comparison test. N/A, no 7 days old larva available.

**Figure 1—figure supplement 2.**
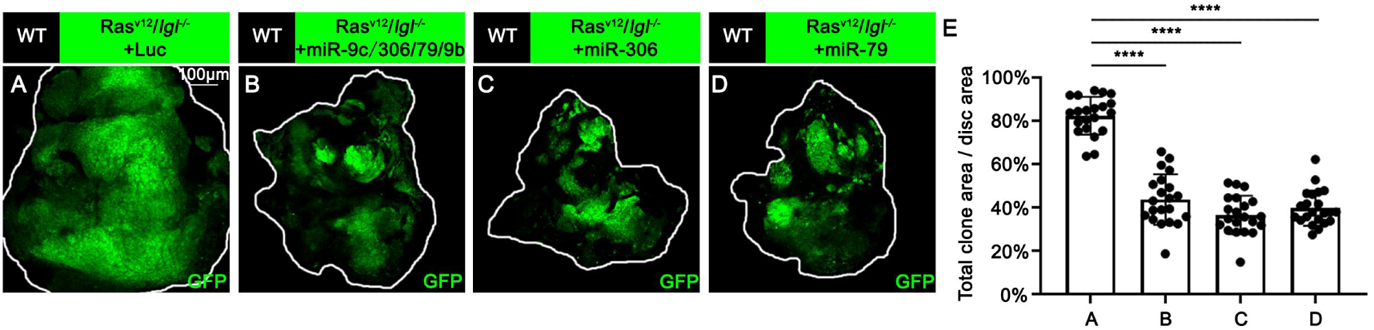
miR-306 and miR-79 suppress Ras^V12^/*lgl^−/−^* tumor growth. (A-D) Eye-antennal disc bearing GFP-labeled clones of indicated genotypes (7 days after egg laying). (E) Quantification of clone size (% of total clone area per disc area in eye-antennal disc) of (A-D). Error bars, SD; ****, p<0.0001 by one-way ANOVA multiple comparison test.

**Figure 1—Source data 1. Source data for Figure 1.**

**Figure 1—figure supplement 1—source data 1. Source data for Figure 1—figure supplement 1**

**Figure 1—figure supplement 2—source data 1. Source data for Figure 1—figure supplement 2**

**Figure 2—figure supplement 1.**
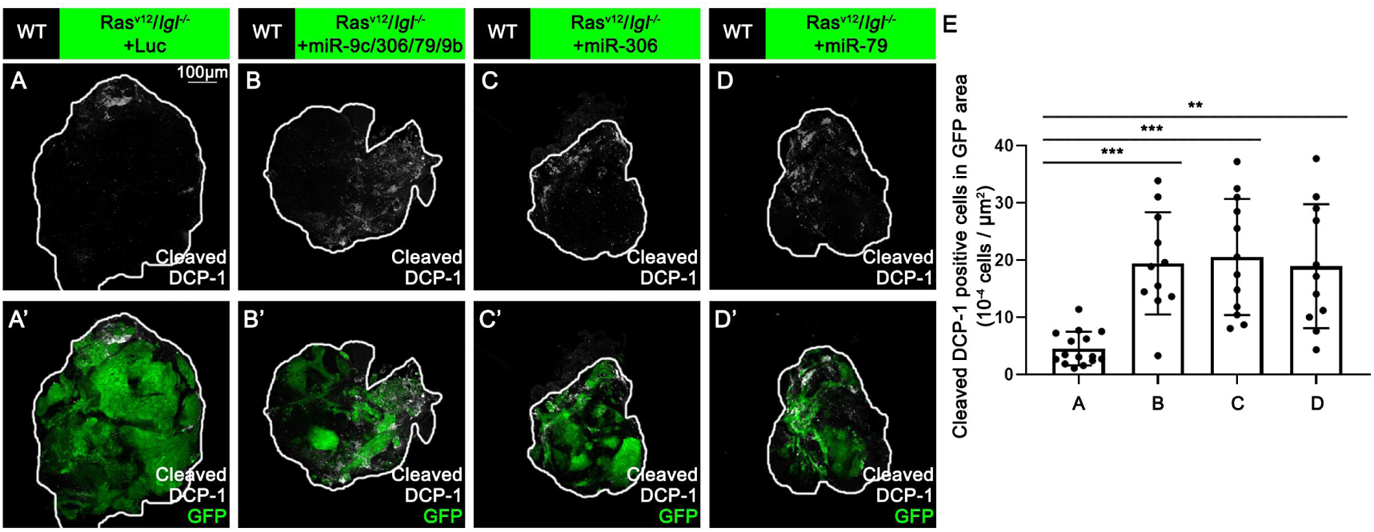
miR-306 and miR-79 induce apoptosis in Ras^V12^/*lgl^−/−^* tumors. (A-D) Eye-antennal disc bearing GFP-labeled clones of indicated genotypes stained with anti-cleaved Dcp-1 antibody (7 days after egg laying). (E) Quantification of dying cells in GFP positive clone area in (A-D). Error bars, SD; ***, p<0.001, ****, p<0.0001 by one-way ANOVA multiple comparison test.

**Figure 2—Source data 1. Source data for Figure 2.**

**Figure 2—figure supplement 1—source data 1. Source data for Figure 2—figure supplement 1**

**Figure 3—figure supplement 1.**
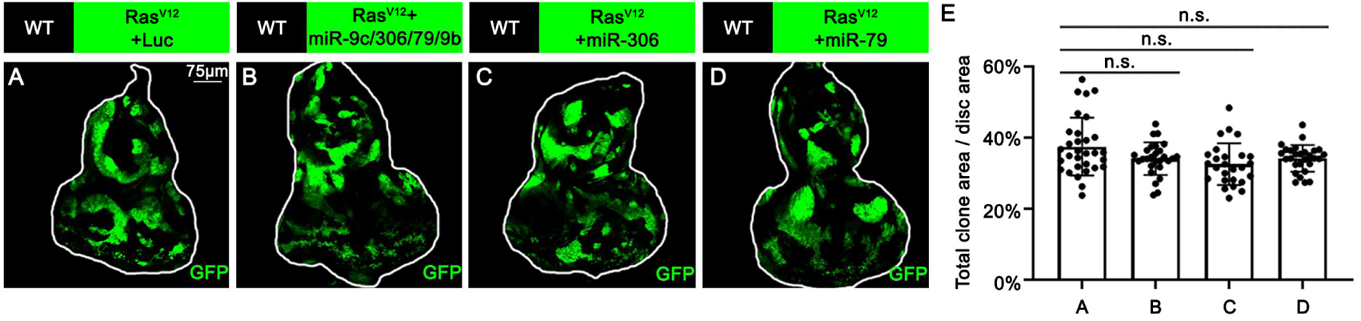
miR-306 and miR-79 do not suppresses Ras^V12^ tumor growth. (A-D) Eye-antennal disc bearing GFP-labeled clones of indicated genotypes (7 days after egg laying). (E) Quantification of clone size (% of total clone area per disc area in eye-antennal disc) of (A-D). Error bars, SD; n.s., p>0.05 by one-way ANOVA multiple comparison test.

**Figure 3—figure supplement 2.**
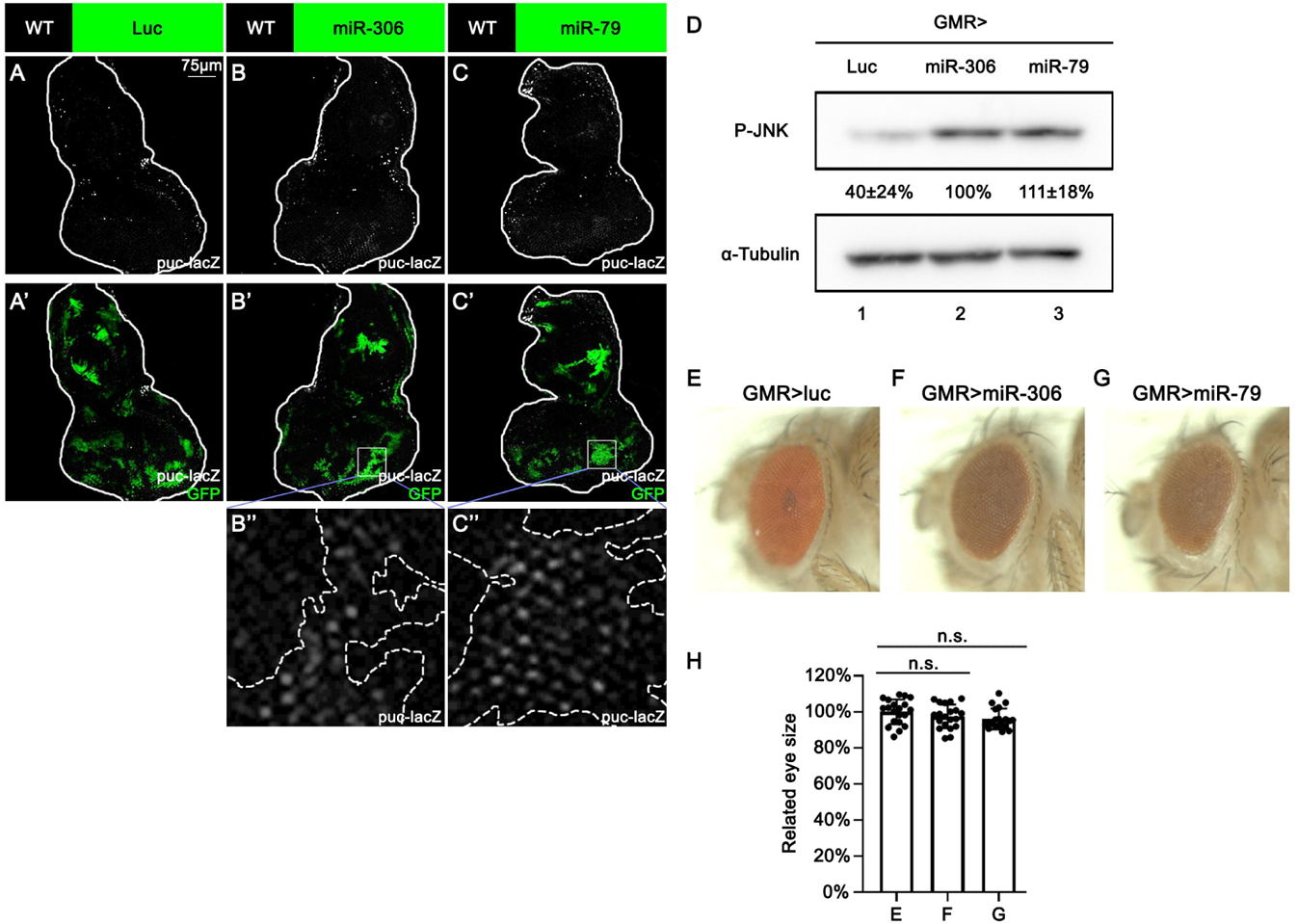
miR-306 and miR-79 promote JNK signaling in the eye-antennal disc and adult eye. (A-C) Eye-antennal disc of indicated genotypes with puc-lacZ background stained with anti-β-galactosidase antibody (5 days after egg laying). (D) Lysates of adult head of indicated genotypes were subjected to Western blots using indicated antibodies. The relative levels of P-JNK protein were presented as means ± SD (normalized to lane 2) from three independent experiments. (E-G) Adult eye phenotype of flies with indicated genotypes. (H) Quantification of adult eye size (normalized to control) of (E-G). Error bars, SD; n.s., p>0.05 by one-way ANOVA multiple comparison test.

**Figure 3—figure supplement 3.**
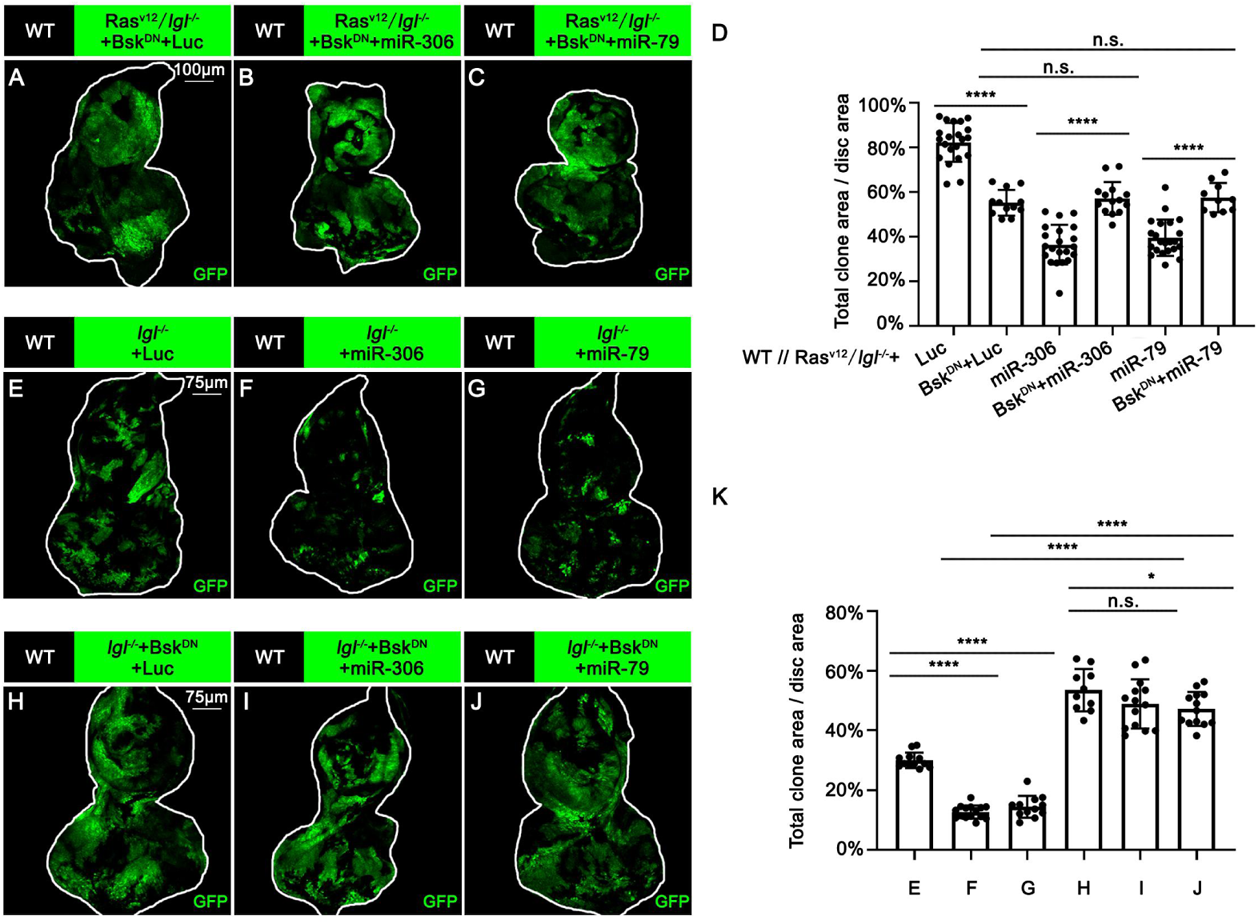
miR-306 and miR-79 suppress Ras^V12^/*lgl^−/−^* tumor growth by promoting JNK signaling. (A-C) Eye-antennal disc bearing GFP-labeled clones of indicated genotypes (7 days after egg laying). (D) Quantification of clone size (% of total clone area per disc area in eye-antennal disc) of indicated genotypes. Error bars, SD; n.s., p>0.05 (not significant), ****, p<0.0001 by two-tailed student’s t test. (E-J) Eye-antennal disc bearing GFP-labeled clones of indicated genotypes (5 days after egg laying). (K) Quantification of clone size (% of total clone area per disc area in eye-antennal disc) of (E-J). Error bars, SD; n.s., p>0.05 (not significant), *, p<0.05, ****, p<0.0001 by two-tailed student’s t test.

**Figure 3—Source data 1. Source data for Figure 3.**

**Figure 3—figure supplement 1—source data 1. Source data for Figure 3—figure supplement 1**

**Figure 3—figure supplement 2—source data 1. Source data for Figure 3—figure supplement 2**

**Figure 3—figure supplement 3—source data 1. Source data for Figure 3—figure supplement 3**

**Figure 4—figure supplement 1.**
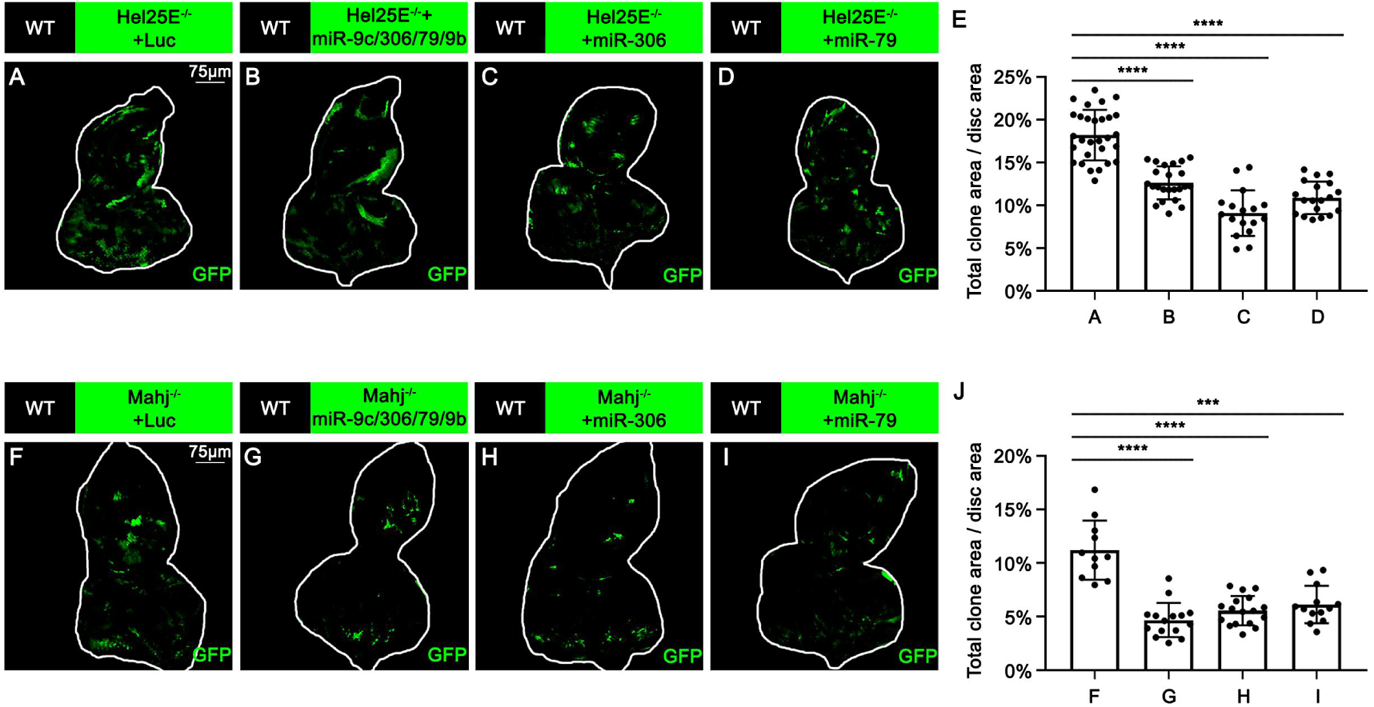
miR-306 and miR-79 promote multiple types of cell competition. (A-D) Eye-antennal disc bearing GFP-labeled clones of indicated genotypes (5 days after egg laying). (E) Quantification of clone size (% of total clone area per disc area in eye-antennal disc) of (A-D). Error bars, SD; ****, p<0.0001 by one-way ANOVA multiple comparison test. (F-I) Eye-antennal disc bearing GFP-labeled clones of indicated genotypes (5 days after egg laying). (J) Quantification of clone size (% of total clone area per disc area in eye-antennal disc) of (F-I). Error bars, SD; ***,p<0.001, ****, p<0.0001 by one-way ANOVA multiple comparison test.

**Figure 4—Source data 1. Source data for Figure 4.**

**Figure 4—figure supplement 1—source data 1. Source data for Figure 4—figure supplement 1**

**Figure 5—figure supplement 1.**
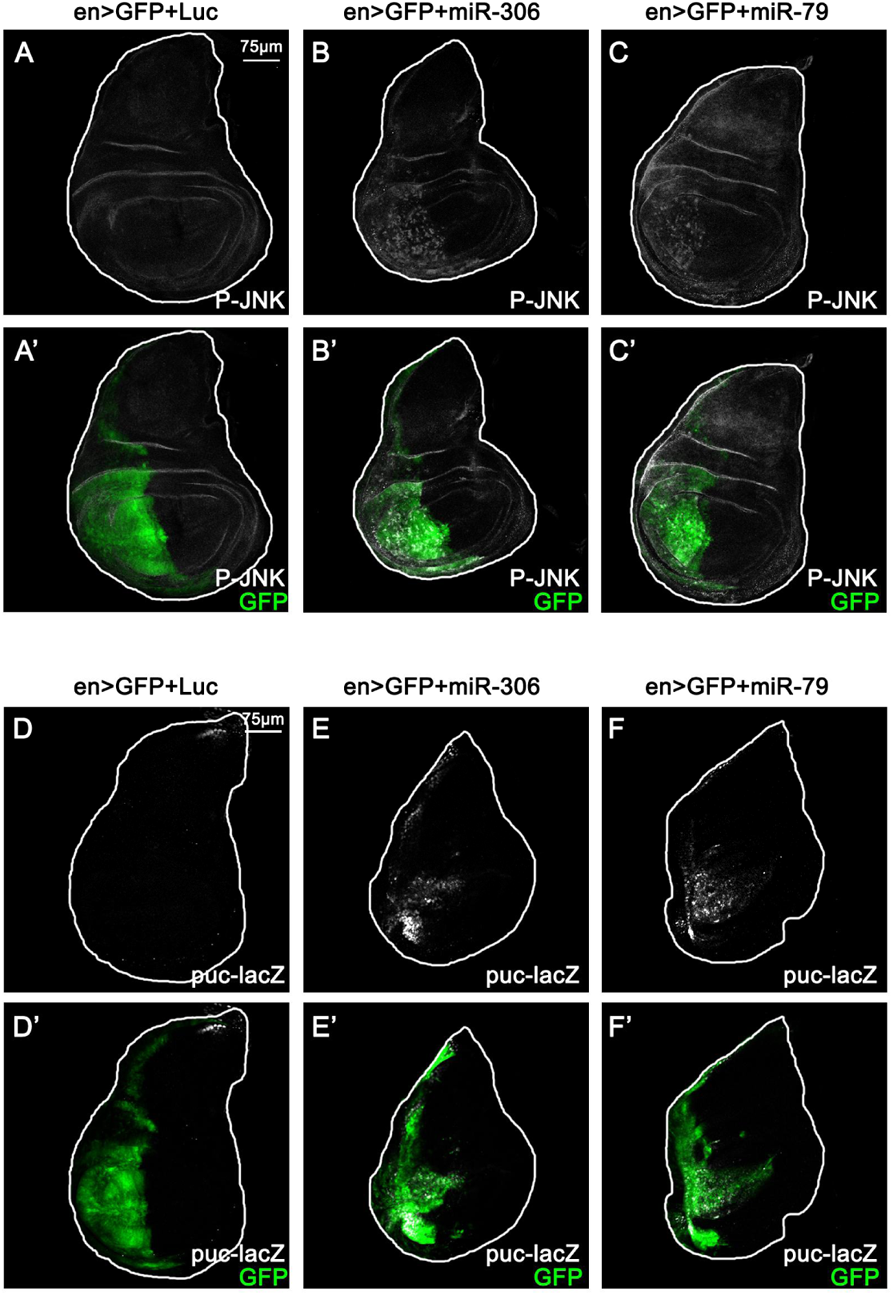
miR-306 and miR-79 promote JNK signaling in the wing disc. (A-C) Wing disc of indicated genotypes stained with anti-Phospho-JNK antibody (5 or 6 days after egg laying). (D-F) Wing disc of indicated genotypes with puc-lacZ background stained with anti-β-galactosidase antibody (5 or 6 days after egg laying).

**Figure 5—figure supplement 2.**
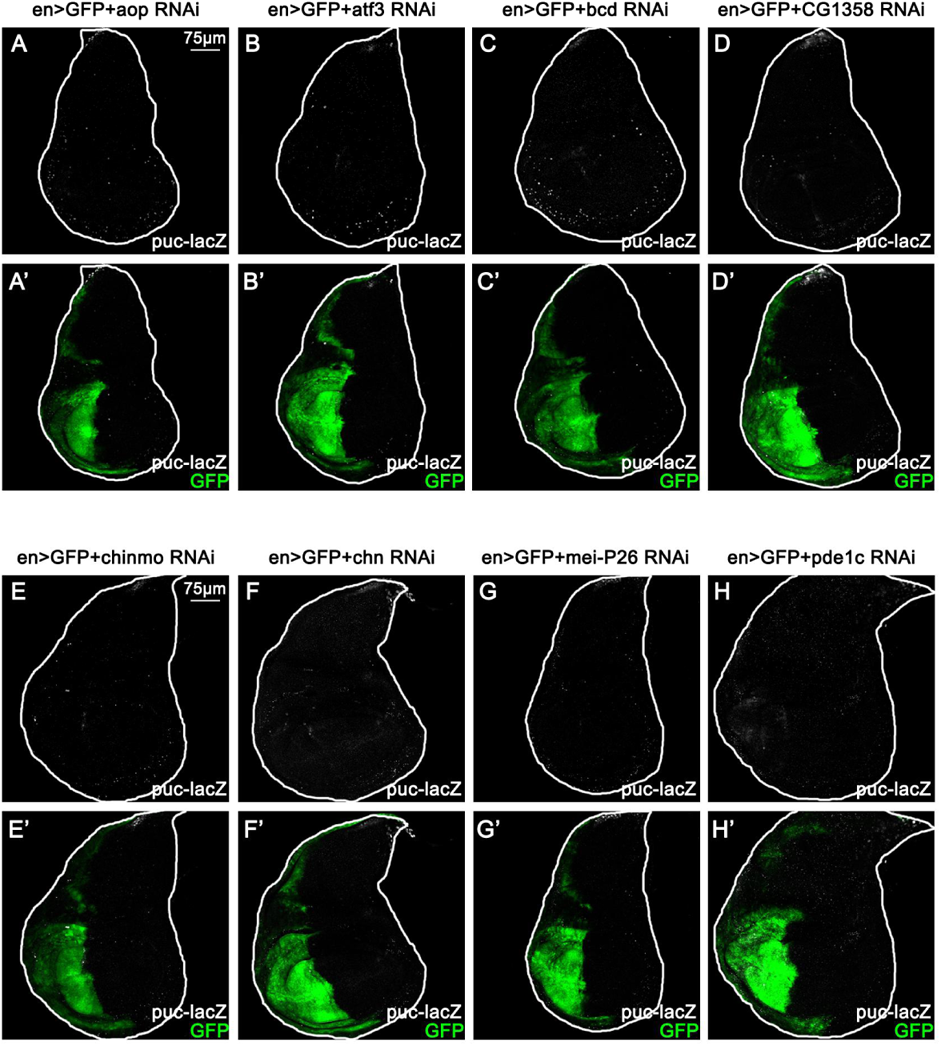
RNAis that target 8 candidate genes do not induce JNK activation in the wing disc. (A-H) Wing disc of indicated genotypes with puc-lacZ background stained with anti-β-galactosidase antibody (5 days after egg laying).

**Figure 5—figure supplement 3.**
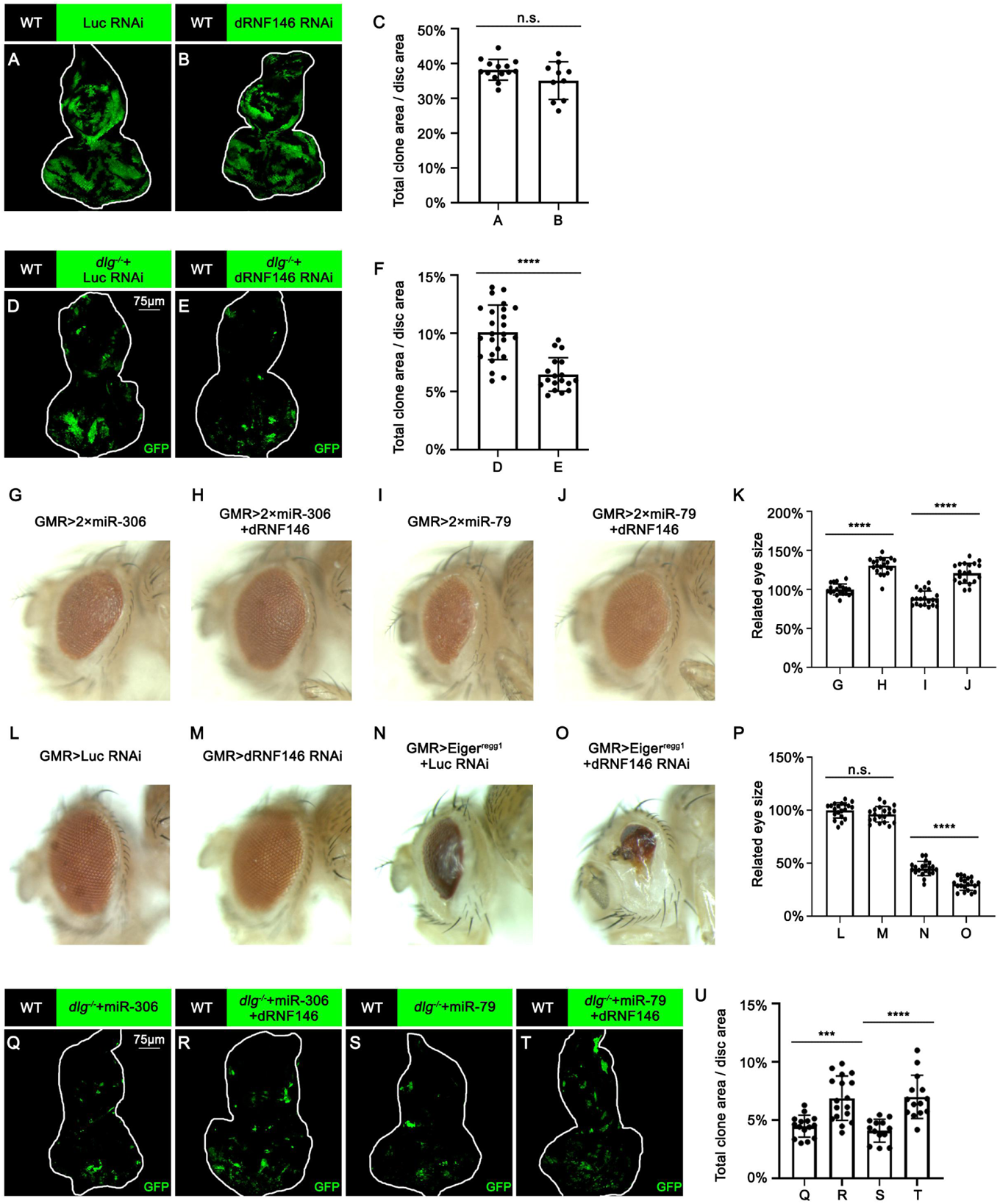
miR-306 and miR-79 promote cell competition by targeting dRNF146. (A-B) Eye-antennal disc bearing GFP-labeled clones of indicated genotypes (5 days after egg laying). (C) Quantification of clone size (% of total clone area per disc area in eye-antennal disc) of (A-B). Error bars, SD; n.s., p>0.05 (not significant) by two-tailed student’s t test. (D-E) Eye-antennal disc bearing GFP-labeled clones of indicated genotypes (5 days after egg laying). (F) Quantification of clone size (% of total clone area per disc area in eye-antennal disc) of (D-E). Error bars, SD; ****, p<0.0001 by two-tailed student’s t test. (G-J) Adult eye phenotype of flies with indicated genotypes. (K) Quantification of adult eye size (normalized to control) of (G-J). Error bars, SD; ****, p<0.0001 by two-tailed student’s t test. (L-O) Adult eye phenotype of flies with indicated genotypes. (P) Quantification of adult eye size (normalized to control) of (L-O). Error bars, SD; n.s., p>0.05 (not significant), ****, p<0.0001 by two-tailed student’s t test. (Q-T) Eye-antennal disc bearing GFP-labeled clones of indicated genotypes (5 days after egg laying). (U) Quantification of clone size (% of total clone area per disc area in eye-antennal disc) of (Q-T). Error bars, SD; ***, p<0.001, ****, p<0.0001 by two-tailed student’s t test.

**Figure 5—figure supplement 4.**
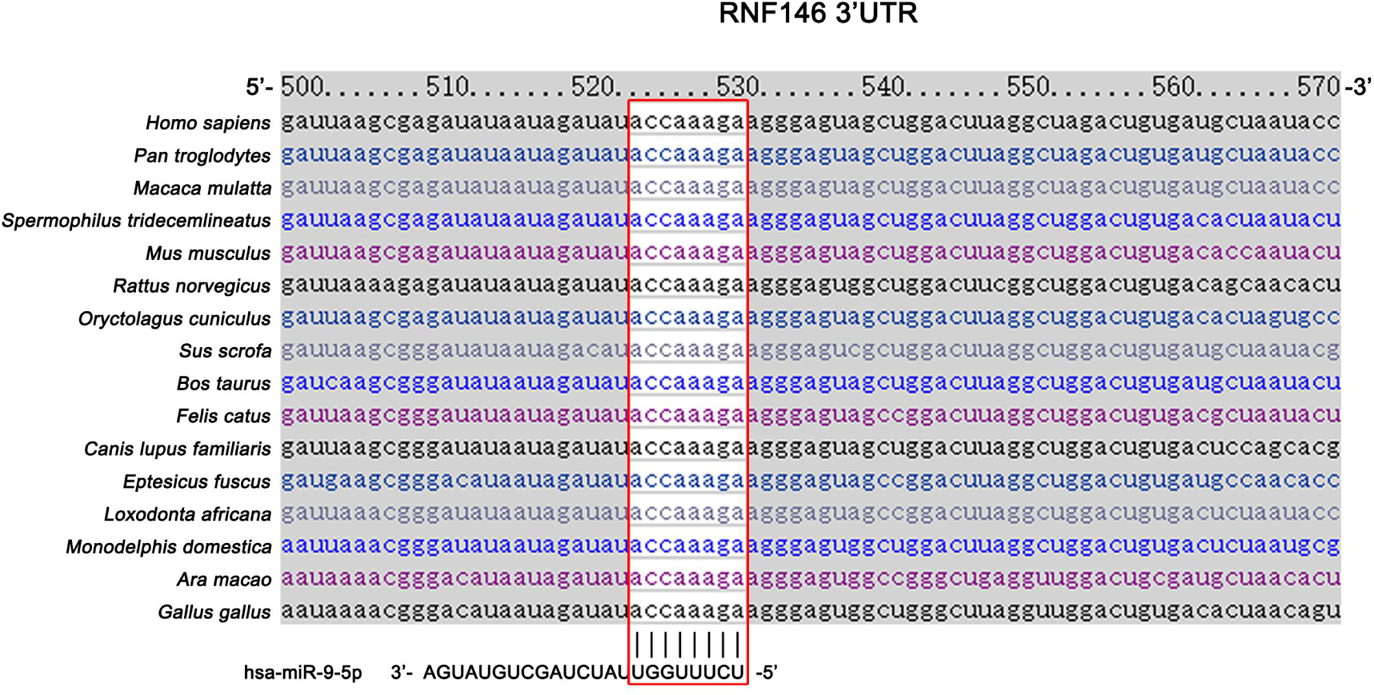
miR-9 is predicted to target mammalian RNF146. Schematic of the miRNA binding sites for miR-9. Red box shows the seed sequence pairing region.

**Figure 5 - Source data 1. Source data for Figure 5.**

**Figure 5—figure supplement 3—source data 1. Source data for Figure 5—figure supplement 3**

**Figure 6 - Source data 1. Source data for Figure 6.**

